# Expanded olfactory system in fishes capable of terrestrial exploration

**DOI:** 10.1101/2022.12.05.518831

**Authors:** Demian Burguera, Francesco Dionigi, Kristina Kverková, Sylke Winkler, Thomas Brown, Martin Pippel, Yicheng Zhang, Maxwell Shafer, Annika L.A. Nichols, Eugene Myers, Pavel Němec, Zuzana Musilova

## Abstract

Smell abilities differ greatly among vertebrate species, as reflected by the exceptional size variability of gene families and brain regions involved in odour detection. However, key environmental factors shaping the molecular and phenotypic evolution of the olfactory sensory system remain mostly unknown. Here, we investigate the association between diverse ecological traits and the number of olfactory chemoreceptors in the genomes of more than two hundred ray-finned fishes. We found independent expansions producing large gene repertoires in nocturnal amphibious fishes, generally able to perform active terrestrial exploration. Furthermore, we reinforced this finding with an additional analysis of a clariid species (*Channallabes apus*), a fish lineage with chemosensory-based aerial orientation. Importantly, we also detected an augmented information-processing capacity in the olfactory bulb of examined nocturnal amphibious species. Overall, we report a convergent magnification of the olfactory system in amphibious fishes potentially enhancing terrestrial orientation, revealing the likelihood of an analogous event in tetrapod ancestors during the water-to-land transition.

## Introduction

At the onset of sensory system evolution in early vertebrates, a few types of chemoreceptor genes acquired roles in the perception of smell^1^. As a result, olfaction in this lineage is mediated by four distinct receptor families: odorant receptors (ORs), trace-amine associated receptors (TAARs), vomeronasal type 1 receptors (V1Rs) and vomeronasal type 2 receptors (V2Rs). Earlier comparative studies on these olfactory (OLF) genes reported striking differences in the number of duplicates among species^2–4^. In particular, studied fishes showed a smaller number of receptors compared to many terrestrial vertebrates, supporting the idea that aerial and water mediums present distinct chemoreception requirements^5^. Nevertheless, high repertoire variability has been recently detected in teleosts^6,7^, although the selective pressures explaining these differences are mostly unknown.

Unlike tetrapods, all OLF gene families are transcribed in a single paired olfactory organ in fishes^8^. This structure, usually organized in a rosette-like epithelium, contains numerous sensory neurons dedicated to the perception of chemical signals. Each of those neurons is specialized in the detection of particular odorant molecules through transcriptional activation of only one among all receptor genes^9^. Moreover, neurons expressing an identical receptor project their axons to a particular region within the olfactory bulb (OB), establishing synapses with the mitral cells located in the same glomeruli^10^. This neural organization allows the decoding of a complex chemosensory input into topological information in the brain. Interestingly, great variation in the relative size of the OB is observed among vertebrate species including fishes, which has been often interpreted as a proxy for smell abilities^11,12^. However, whether genetic variability involving OLF gene repertoires is linked to the information-processing capacity in the brain, reflecting a functional impact, remains largely unexplored.

### Independent olfactory gene expansions in ray-finned fishes

We investigated the evolutionary dynamics of the OLF genes in a large and diverse set of ray-finned fish species (Supplementary Table 1). We identified OR, TAAR, V1R and V2R genes in high-quality genomes of more than two hundred species from approximately fifty taxonomic orders (Supplementary Fig. 1a). While half of the species presented less than 250 OLF genes in total, we found a striking variability ranging from 29 genes in the common seadragon to more than 1300 in the ropefish (Fig. 1a). Remarkably, we observed large repertoires with more than twice the median value (>500 OLF genes) in species from seven distinct orders (Fig. 1a).

**Fig. 1:**
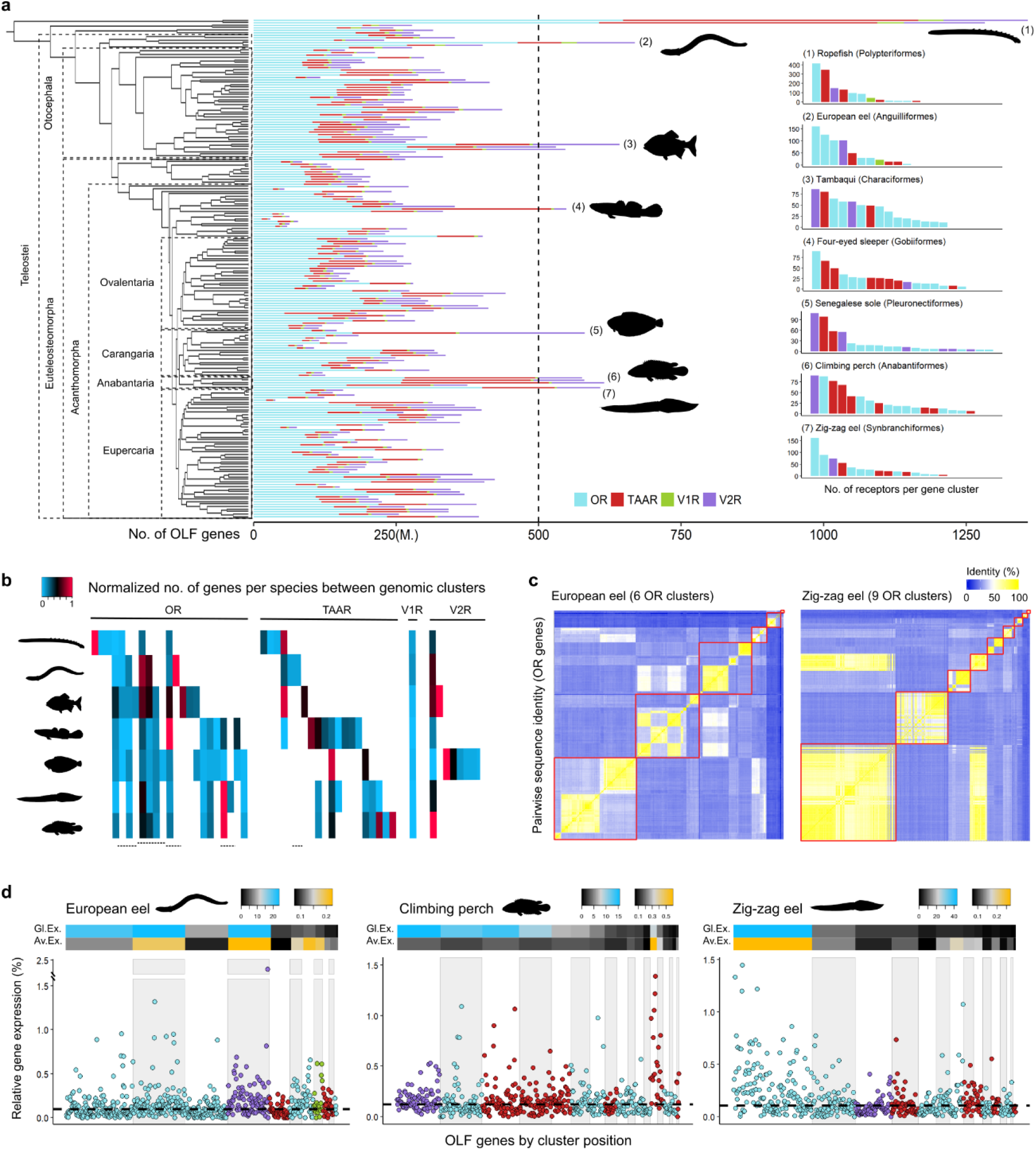
Genomic and transcriptomic dynamics of expanded OLF gene clusters. Loci containing less than five receptor genes are excluded for visualization purposes. **(a)** Total number of OLF genes in 201 ray-finned fish species according to gene family. Median value is indicated (M.). Right, species with higher number of genes from those order having more than 500 OLF genes. **(b)** Normalized number of receptor genes in each genomic cluster per species. Homologous gene clusters (cells) are displayed in columns. White cells represent absence of homologous gene clusters. Dashed lines indicate clusters of homologous origin separated by genomic rearrangements. Clusters containing the highest number of genes in distinct species are mostly non-coincident **(c)** Heatmaps showing sequence identity between individual OR receptors in the European eel and zig-zag eel. Gene clusters delimited by red squares. **(d)** Relative gene expression levels of individual receptors in three selected species displayed in genomic order within clusters. Gene families are coloured as in panel a. Median value indicated by dashed line. Genomic clusters are highlighted in white and light grey alternatively. On top, heatmaps showing the percentage of global accumulated OLF expression (Gl. Ex.) and average per gene expression (Av. Ex) of each gene cluster.

Lineages presenting such expansions include bichirs, true eels, characins, sleeper gobies, true soles, labyrinth fishes and spiny eels. In particular, the total number of receptor genes in the bichirs is comparable to many olfactory-oriented mammals^13^. Contrary to other clades, none of these species experienced lineage-specific whole genome duplications (Supplementary Fig. 1b), suggesting that olfactory gene dynamics are fast and probably associated with macroevolutionary ecological drifts.

As OLF genes are organized in clusters, we investigated the genomic architecture of the detected expansions. We found the number of gene clusters variable among distantly related clades due to gains, losses and genomic rearrangements (Fig. 1b). In species with an expanded repertoire, we observed an uneven distribution of genes per cluster, with a few loci accumulating a high number of receptors (Fig. 1a). These local inflations affected OR, TAAR and/or V2R families unequally in each lineage, producing specific amplification patterns affecting distinct gene clusters (Fig. 1b). Because expansions are generated by tandem duplications, groups of receptors from the same gene clusters present the highest sequence similarities within species (Fig. 1c). As an exception, we identified an OR cluster in the zig-zag eel containing receptors very similar to genes from a remote cluster, suggesting a very recent evolutionary origin. However, identical receptors are generally scarce, indicating the prevalence of coding divergence after duplication events (Supplementary Fig. 1c).

To study how gene number evolution affects the transcriptional balance between receptors in the olfactory organ, we sequenced and analysed the transcriptomes of eight species with differently sized repertoires. We found substantial variation in the transcriptomic abundance of OR, TAAR, V1R and V2R families among lineages, roughly matching their genomic proportions (Supplementary Fig. 2a-b). Moreover, we observed a minority of individual receptors (15-20%) accumulating half of the overall OLF gene expression in studied fishes (Supplementary Fig. 2c). To investigate whether expanded gene clusters are represented at the transcriptomic level, we focused in more detail on the three species with greater repertoires (>600 genes) (Fig. 1d). We found the largest gene cluster in the zig-zag eel as the most abundant both in terms of accumulated and average gene expression. Similarly, the other fishes also concentrate higher fractions of the total OLF expression in their bigger clusters. However, these two species showed higher transcriptional rates per gene in clusters other than the largest, with the remarkable case of a small but extremely expressed TAAR cluster in the climbing perch. Overall, transcriptomic abundances at the family and cluster level globally reflect gene expansions, despite regulatory mechanisms able to locally enhance the expression of individual receptors in a locus-specific manner.

### Olfactory molecular evolution linked to ecology and brain processing capacity

To identify environmental causes influencing the number of OLF genes in ray-finned fishes, we estimated the effect of distinct ecological, life-history and behavioural traits with a potential impact on smell evolution. We found a significant deviation towards larger repertoires in three of the studied factors: freshwater habitats, nocturnal activity, and amphibious behaviour (Fig. 2a). Furthermore, we observed that fishes presenting the exact combination of these three traits include many of the more expanded repertoires in our data set (Fig. 2c). Interestingly, most of these species are reported to perform active terrestrial exploration to find new water bodies or prey^14,15^. Indeed, testing the influence of terrestrial exploration on the number of receptors as a single trait revealed the largest significant effect detected with our model (Fig. 2b). In contrast, diurnal amphibious fishes such as mudskippers, bullheads or some toothcarps, rarely found more than a few meters from water^16^, do not present enlarged olfactory repertoires (Fig. 2d). Perhaps relatedly, some of these diurnal species are described to use vision to orientate towards their original aquatic environment^17,18^. Thus, we suggest that nocturnal amphibious lineages might particularly benefit from an increase of their OLF gene repertoires, especially for terrestrial exploratory excursions involving relatively large distances.

**Fig. 2:**
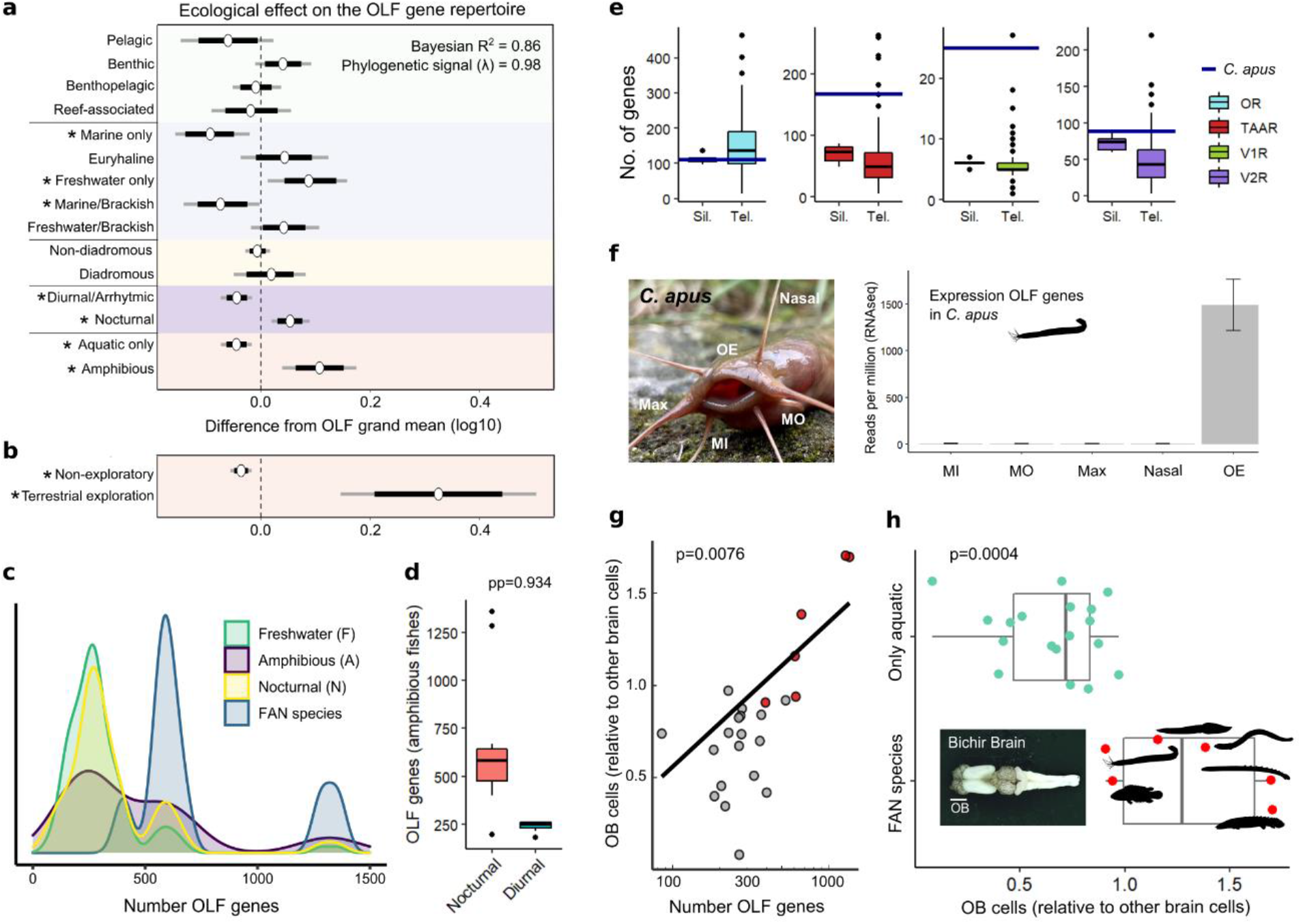
Ecological links to olfactory system evolution in ray-finned fishes. **(a,b)** Statistical effect of several ecological traits on the number of OLF genes, with highest density intervals (HDI) of 80% (black) and 95% (grey). Colours delimit related factors containing exclusive sets of species. Statistical significance (95% HDI does not include 0) is marked with an asterisk. **(c)** Density plot with the number of OLF genes in non-exclusive ecological groups (freshwater, n=125; amphibious, n=19; nocturnal, n=62), plus a subset of fishes presenting all the above traits (FAN, n=9). **(d)** Differences in the OLF repertoire size between nocturnal (n=11) and diurnal (n=8) amphibious fishes. Posterior probability (pp) of nocturnal being higher than diurnal is indicated. **(e)** Number of genes for each receptor family identified in our genome assembly of eel catfish (*C. apus*), compared to other Siluriformes (Sil; n=9) and teleost (Tel; n=195). **(f)** Left, picture of *C. apus* showing position of the olfactory epithelium (OE) and the four types of barbels: nasal, maxillary (Max), mandibular outer (MO) and mandibular inner (MI). Right, number of RNA-seq reads (per million reads) from barbels and OE mapped to annotated OLF genes in *C. apus* (SD shown). **(g)** PGLS model showing a significant correlation between the number of OLF genes and the covariance in the number of OB and other brain cells in twenty-four species (x-axis is log-transformed). FAN species coloured in red. **(h)** Significantly higher proportion of OB cells in FAN species (n=6) compared to the rest of fishes (n=18) after phylogenetic correction.

Terrestrial orientation through aerial chemoreception in fish has been recently reported for the first time in a clariid species^19^. This siluriform clade is known to emerge from water nocturnally to move between ponds or streams. In order to investigate whether this amphibious lineage also presents an expanded OLF gene repertoire, we sequenced the genome of the eel catfish, *Channallabes apus (C. apus)* (see Methods), a tropical clariid able to hunt for terrestrial prey^20,21^. Using our high-quality assembly, we identified expansions of the TAAR and V1R gene families compared to other siluriform and teleost species (Fig. 2e), reinforcing the association of nocturnal amphibious behaviour with increased receptor repertoires. It is worth mentioning that aerial chemoreception in clariids was previously hypothesized to function mainly through the barbels^19^, a gustatory organ. In particular, it was proposed that the nasal barbels might have co-opted the expression of olfactory receptors to increase their sensory capabilities. However, we only detected robust expression of OLF genes in the olfactory epithelium and not in the barbels of *C. apus*, indicating that the sensory role of these receptor families is restricted to smell and not taste (Fig. 2f).

We next studied whether fishes with large OLF gene repertoires present an augmented information-processing capacity in their olfactory bulbs (OB), frequently used as a proxy for smell functional abilities^12^. Nevertheless, comparison of volume ratios in distantly related species are problematic as similarly sized brains can differ in the number and distribution of neurons^22^. Therefore, we directly estimated the cell number of dissected brain parts from twenty-four ray-finned fish species using the isotropic fractionator (see Methods). With this approach, we detected a positive correlation between the amount of receptor genes and the number of OB cells relative to other brain cells (PGLS, p=0.076, Fig. 2g). Furthermore, nocturnal amphibious fishes presented a higher proportion of OB cells compared to the rest of species, after correcting for phylogeny and brain size (p= 0.0004, Fig. 2h). Hence, our results link the detected genetic variability with a sensory phenotypic trait and provide further evidence for enhanced olfactory abilities in amphibious lineages with expanded OLF gene repertoires.

Last, we investigated whether gene expansions in nocturnal amphibious fishes occurred in the same subtype of receptors most abundant in tetrapods, called γ-OR genes^5^ (Supplementary Fig. 3). While we found them absent or scarce in amphibious teleost species, we identified a substantial number of γ-OR in the bichirs (Fig. 3). Interestingly, we observed the majority of γ receptors in this lineage located in a gene cluster that was lost before the teleost radiation (Fig. 3), but still present in other ray-finned fishes. Thus, increments in the number of chemoreceptors happened in distinct subtypes in most amphibious fishes compared to tetrapods, maybe related to this ancient loss.

**Fig. 3:**
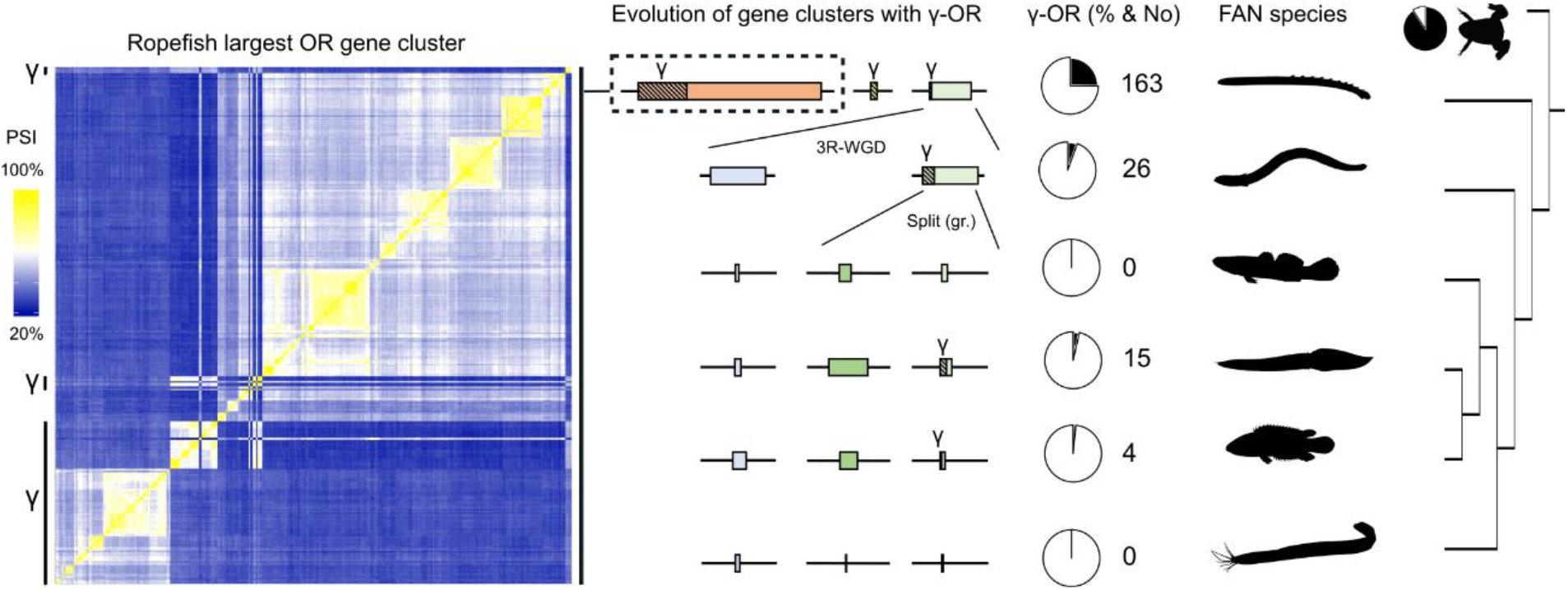
Gamma-OR genes scarcity in amphibious teleost after gene cluster loss. Evolutionary comparison of homologous gene clusters containing γ-OR genes in a subset of FAN species reveal the highest number and proportion of this receptor subtype in bichirs, found in three distinct gene clusters. Analysed teleost species present much smaller γ-OR repertoires, restricted to a unique gene cluster after the loss of γ-OR containing loci. Left, pairwise sequence identity (PSI) at the protein level between receptors from the largest gene OR cluster in the ropefish. Regions containing mostly γ-OR are marked. Middle, boxes represent the relative size of gene clusters homologous to those containing γ-ORs in the bichirs. The proportion of γ-ORs (hatched) is also at scale. The third round of whole genome duplication (3R-WGD) at the base of the teleost clade and a genomic rearrangement (gr.) in clupeocephalans are indicated. The proportion of γ-OR in pie charts is calculated over the complete OR repertoire in each species. From top to bottom, species in the figure include the ropefish (Polypteriformes), European eel (Anguilliformes), four-eyed sleeper (Gobiiformes), zig-zag eel (Synbranchiformes), climbing perch (Anabantiformes) and *C. apus* (Siluriformes). Dominant γ-OR proportion in an amphibian outgroup is also shown (*Xenopus tropicalis*). Ropefish and *Xenopus* silhouettes were downloaded from http://phylopic.org/.

## Discussion

We explored potential associations of several ecological and behavioural traits with olfaction-related changes in ray-finned fishes. We found that nocturnal amphibious lineages have convergently expanded their olfactory sensory system molecularly and in terms of OB processing capacity in the brain. From a greater phylogenetic perspective, this finding resembles the case of amphibians, which present particularly high numbers of OLF genes presumably linked to distinct olfaction requirements in their aquatic and terrestrial forms^23^. On the other hand, cetaceans and sea snakes have severely reduced OR repertoires compared to land species^24^, showing a consistent evolutionary response to the physical medium in both directions in vertebrates. Therefore, we suggest that these increments in the odour processing capacity of nocturnal amphibious fishes are likely connected to facilitate terrestrial orientation, a potentially useful skill during the search of new water bodies or prey. Moreover, we further strengthened this evolutionary scenario with on purpose genome sequencing and analysis of a clariid species (*C. apus*), an amphibious fish clade with previously described aerial chemosensory abilities^19^. Nonetheless, as we also identified substantial gene repertoires in a small subset of exclusively aquatic species, additional ecological causes leading to olfactory genomic variability in fishes remain to be discovered.

In addition, we also identified in the bichirs a substantial number of genes homologous to the most abundant receptor subtype in land vertebrates, named γ-OR. This particular group of chemoreceptors was considered to exclusively detect airborne odorants by earlier studies based on their absence in the first studied teleost genomes^5^. Although not completely missing, we hypothesize that the prevailing scarcity of γ-OR genes in this lineage might be at least partially related to the loss of a particular genomic cluster before the diversification of teleosts. However, this receptor subtype is markedly present in non-amphibious fishes from other clades such as the spotted gar or the coelacanth^25^, suggesting an ancestral function in water environments. In fact, it is increasingly recognised that aquatic animals are able to detect small hydrophobic and volatile compounds^25,26^. Thus, we speculate that some of the chemosensory expansions found in amphibious species might occurred in receptor genes already able to bind volatile molecules present in water. Finally, given that extant sarcopterygian fishes show modest OLF gene repertoires^27^, we propose that an analogous magnification of the olfactory system probably happened during the first steps of the water-to-land transition in early tetrapods. Thereby, fast chemosensory changes could have preceded many key morphological adaptations in arguably one of the most crucial events during vertebrate evolution.

## Methods

### OLF genes annotation and analysis of clusters

We developed a pipeline to identify olfactory genes in 201 high-quality ray-finned fish with publicly available genome assemblies (Supplementary Table 1). First, available reference protein sequences of OR, TAAR, V1R and V2R receptors from several species belonging to separate actinopterygian clades (spotted gar, zebrafish, stickleback, fugu, tongue sole, European seabass, Nile tilapia, mandarin fish, salmon and medaka)^2–5^ were blasted against target genome assemblies (tblastn, e-value < 1e-35). Only best hits were recovered in case of overlapping between queries. We built preliminary gene models grouping high scoring pairs belonging to different parts of the same queries to account for putative introns when necessary (maximum intron size of 10Kb for OR, TAAR and V1R and 30Kb for V2R). We extracted the genomic sequence spanning the gene models with an extension of three kilobases on each side. These sequences were processed with Exonerate (EMBL-EBI) using the original reference to generate optimized gene models (minimum identity of 35%). Next, we obtained the coding sequence of these new models with TransDecoder (https://github.com/TransDecoder/TransDecoder) and kept those equal or larger than 275 aminoacids for OR, TAAR, V1R, and 700 aminoacids for V2R. The obtained sequences were blasted against a database containing OR, TAAR, V1R or V2R protein sequences and other G-protein coupled receptor families (HODER database) to filter out non-OLF receptors (blastp, e-value < 1e-35). We repeated the whole process in each species using the new obtained gene models as references to further recover divergent unannotated OLF genes (Supplementary Data 1).

Also, to avoid wrong estimates in our analysis, we restricted the dataset to publicly available high-quality genome assemblies. We only collected those assemblies that surpassed two *a priori* and one *a posteriori* filters: (i) The N50 metric divided by the total length of the genome was at least 10Mb/Gb. (ii) The minimum required BUSCO.v3 completeness (actinopterygii10 database) was 90% for all species except for Polypteriformes, as a minimum of 85% was tolerated for the outgroup lineage within our dataset. (iii) To account for excessive genomic fragmentation, annotated OR genes needed to be distributed in less than twenty scaffolds or chromosomes, while in species with lineage-specific whole genome duplications up to thirty scaffolds were allowed. Gene cluster homologies among species were inferred based on syntenic genes flanking studied loci (Supplementary Table 2). To unveil the cluster duplication dynamics, protein-level identity values from pairwise sequence alignments of analysed receptor genes were calculated with Clustal omega (EMBL-EBI). CD-HIT (https://github.com/weizhongli/cdhit) was used for grouping OLF genes with distinct levels of sequence identity.

### Fish samples

Some of the studied species were obtained through the aquarium trade (Aquarium Glaser GmbH, Rodgau, Germany): *Anabas testudineus, Pygocentrus nattereri, Electrophorus electricus, Erpetoichthys calabaricus, Polypterus senegalus, Mastacembelus armatus, Xenentodon cancila, Thalassophryne amazonica, Gambusia affinis, Xiphophorus helleri* and *Channallabes apus*. Other fish were sampled from wild, semi-wild or captive facilities in the Czech Republic: *Acipenser ruthenus, Anguilla anguilla, Leuciscus idus, Rutilus rutilus, Gobio gobio, Esox lucius, Perca fluviatilis, Sander lucioperca, Oncorhynchus mykiss, Salmo trutta* and *Thymallus thymallus*. Last, a few marine species were obtained from fishing catches in Genoa (Italy): *Gadus morhua, Sparus aurata* and *Dicentrarchus labrax*. Two or three adult individuals from each species were processed for transcriptomic and/or brain analysis based on availability. We obtained all necessary permits for sampling and processing life fish samples. Handling permit was issued by the Ministry of Agriculture of the Czech Republic (CZ03061), and collection permit for this study was issued for all localities by the Czech Fishing Association as officially authorized by the Ministry of Agriculture of the Czech Republic.

### Transcriptomic analyses

We dissected the olfactory organ from two individuals of eight species with diverse OLF gene repertoire sizes: *Anguilla anguilla* (669), *Pygocentrus nattereri* (530), *Electrophorus electricus* (332), *Thymallus thymallus* (85), *Salmo trutta* (183), *Thalassophryne amazonica* (141), *Anabas testudineus* (615) *and Mastacembelus armatus* (608). Total mRNA was extracted with RNeasy Mini Kit (Qiagen). RNA sequencing was performed by Novogene to obtain 30-55 million reads from an Illumina platform (paired end, 150 × 2 base pairs) per sample (Supplementary Table 3). Reads were filtered with Fastp software^28^. We estimated the expression of annotated receptors by balancing the number of uniquely and ambiguously mapped reads using Salmon^29^. As each sensory neuron expresses only one receptor, and mRNA levels are known to correlate with the number of neurons expressing a given receptor^30,31^, we calculated the relative abundance of every transcript among all OLF receptors for interspecific comparison. This approach also reduced potential bias due to contamination of surrounding non-olfactory tissues compared to absolute measurements. Mean values obtained from biological replicates for each gene were used. Nevertheless, one particular V2R gene located outside of the main gene cluster of this family and lacking receptor activity is known to be co-expressed with other V2R genes in previously studied vertebrates^32^. Accordingly, we identified homolog singletons of this gene with very high expression in our studied fish species (data not shown). Hence, we decided to eliminate it from the expression analysis to avoid overestimation in V2R abundance. In the case of *Channallabes apus* analysis, read counts per million sequenced reads were used for direct comparison of the expression level of OLF genes among tissues.

### Time-calibrated phylogenetic tree

We downloaded the protein sequences of 640 exons from zebrafish (Ensembl), selected by a previous study as optimal for phylogenomic analysis in teleost fishes^33^. We blasted these exons against all the genome assemblies in our dataset (blastn, e-value < 1e-50). We retrieved the sequences corresponding to the best hits and blasted them back to the zebrafish genome (GRCz10). Those exons with reciprocal best hit were considered homologous and used in the subsequent steps. Homologous exons were aligned with MAFFT^34^ and the two first and last nucleotides were trimmed from each alignment to avoid potential gaps caused by incomplete sequence retrieval near the splice sites. Exon alignments were concatenated (Supplementary Data S2), and a maximum-likelihood phylogenetic analysis was conducted using IQTREE2^35^, with a topological constrain for non-teleost species following current molecular phylogenies^36^. Last, the obtained tree was processed with BEASTv2.5 to generate a time-calibrated phylogeny (Supplementary Fig. 4).

### Analysis of ecological features

We gathered information from public databases and scientific literature for several ecological parameters with a potential impact on the evolution of smell abilities (Supplementary Table 4). Information involving habitat zone (Pelagic, Benthic, Benthopelagic, Reef-associated), salinity (Marine, Freshwater, Brackish) and diadromous migrations was mainly collected from FishBase using “rfishbase” R package. Amphibious behavior was determined following available studies^14,15^. Data regarding nocturnal, arrhythmic, or diurnal activity was obtained from the literature for approximately 80% of species in our dataset (Supplementary Table 5). Species presenting a nocturnal/crepuscular pattern were classified as nocturnal, while those having a diurnal/crepuscular pattern were considered diurnal in our analysis. To assess the effect of the studied ecological parameters on the size of the OLF gene repertoires, we used Bayesian phylogenetic multilevel models using the “brms” package^37^. The response variable was log10-transformed number of OLF genes and the explanatory variables were the different ecological categories (Supplementary Table 6). We performed robust linear regression using the student family, specifying the following weakly informative priors: prior(normal(0, 1), “b”), prior(normal(2, 0.5), “Intercept”), prior(student_t(3, 0, 1), “sd”), prior(student_t(3, 0, 1), “sigma”), prior(gamma(3, 0.1), “nu”). Prior predictive checks confirmed that these priors set reasonable expectations for the model coefficients. To check for prior sensitivity, we also ran the model with default priors and the model inference was not changed. We ran 4 MCMC chains for 20,000 iterations with a warm-up period of 10,000 and thinning every 10 steps, resulting in 4000 post-warmup draws. Proper chain mixing was checked by visually inspecting the trace plots. Additionally, Rhat for all parameters was < 1.003 and the bulk and tail effective sample sizes for all parameters were at least 500 and typically over 80% of total post-warmup draws, indicative of model convergence. Bayesian R^2^ was calculated using the bayes_R2 function in the brms package. The phylogenetic signal was estimated using the hypothesis function in the brms package, following the brms vignette (https://cran.r-project.org/web/packages/brms/vignettes/brms_phylogenetics.hhtm).

### *Channallabes apus* genome sequencing

Long genomic DNA (gDNA) of *Channallabes apus* was extracted with the circulomics Nanobind Tissue Big DNA kit (part number NB-900-701-01, protocol version Nanobind Tissue Big DNA Kit Handbook v1.0 (11/19)) according to the manufacturer’s instructions. In brief, liver tissue was minced to small slices on a clean and cold surface and finally homogenized with the TissueRuptor II device (Qiagen) making use of its maximal settings. After complete tissue lysis, remaining cell debris were removed, and the gDNA was bound to circulomics Nanobind discs in the presence of Isopropanol. High molecular weight (HMW) gDNA was eluted from the nanobind discs in elution buffer (EB). The integrity of the HMW gDNA was determined by pulse field gel electrophoresis using the Femto Pulse device (Agilent

Technologies) showing a clear peak 127 kb in length. All pipetting steps of ultra-long and long gDNA were done carefully with wide-bore pipette tips.

The HMW gDNA was used as recommended by Pacific Biosciences according to the ‘Guidelines for preparing HiFi SMRTbell libraries using the SMRTbell Express Template Prep Kit 2.0 (PN 101-853-100, version 03). In summary, HMW gDNA was sheared twice with the MegaRuptorTM device (Diagenode) applying the 25 and 20 kb shearing options. 5 ug sheared gDNA went into library preparation. The PacBio SMRTbellTM library was size selected for fragments larger than 5,5 kb with the BluePippinTM device according to the manufacturer’s instructions and resulted in a PacBio HiFi library of 11,8 kb in size (Fragment Analyzer, Agilent Technologies). The size selected library ran on one Sequel II SMRT cells with the SEQUEL II sequencing kit 2.0 for 30 hours on the SEQUEL II.

Chromatin conformation capturing of *Channallabes apus* chromatin was performed with the ARIMA HiC+ Kit (Material Nr. A410110) following the user guide for animal tissues (ARIMA-HiC 2.0 kit Document Nr: A160162 v00). In brief, circa 50 mg of flash-frozen powdered tissue were chemically crosslinked in enriched nuclei. The crosslinked genomic DNA was digested with the restriction enzyme cocktail consisting of four restriction enzymes. The 5’-overhangs were filled in and labelled with biotin. Spatially proximal digested DNA ends were ligated. The ligated biotin containing fragments were enriched and used for Illumina library preparation, which followed the ARIMA user guide for Library preparation with the Kapa Hyper Prep kit (ARIMA Document Part Number A160139 v00). The barcoded HiC library run on an S4 flow cell of a NovaSeq6000 with 2x 150 cycles.

### *Channallabes apus* genome assembly

We created PacBio CCS reads (read quality > 0.99) from the *Channallabes apus* subreads.bam file using PacBio’s ccs command line tool (version 6.3.0). We obtained 28.36 Gb high quality CCS reads (HiFi reads) with a N50 of 11.58 Kb. To further increase the read quality and coverage we applied the tool DeepConsensus (v0.2, on PacBio reads within 98.0-99.5% read accuracy)^38^ and gained an overall yield of 30.02 Gb (N50: 11.63 Kb). PacBio reads containing the PacBio adapter sequence were filtered out by applying a blast^39^ search providing the PacBio adapter sequence and the following arguments “reward 1 -penalty -5 -gapopen 3 -gapextend 3-dust no -soft_masking false -evalue 700 -searchsp 1750000000000 -outfmt 7”. Initial contigs were generated using HiFiasm (v0.16.1-r375)^40^ with parameters --primary -l0 and alternative haplotigs were purged using purge-dups (v1.2.3)^41^. To create the set of alternative contigs, purge-dups was also run on the alt assembly from hifiasm combined with the purged output of running purge-dups on the primary contigs. Initial scaffolding of the primary assembly was performed by mapping HiC reads to the primary contigs using bwa-mem (v0.7.17-r1198-dirty) and mappings were filtered following the VGP arima mapping pipeline (https://github.com/VGP/vgp-assembly/tree/master/pipeline/salsa). The final bed file was given to yahs^42^ for scaffolding. Scaffolds were then manually curated into chromosomes using higlass (v2.1.11) to visualise the HiC data. Finally, the primary and alt assemblies were polished by mapping the HiFi reads to the assemblies using pbmm2 (v1.3.0) [https://github.com/PacificBiosciences/pbmm2.git] with arguments --preset CCS -N 1 and variants called using deepvariant (v0.2.0) with --model_type=PACBIO. Finally, errors were corrected in the assembly by filtering the vcf file given by deepvariant (v0.2.0)^43^ with bcftools view (v1.12)^44^ with arguments -i ‘FILTER=\”PASS\” && GT=\”1/1\”‘ and a consensus called with bcftools consensus. Finally, we obtained a principal genome assembly of approximately 1,16 Gb in size, 102 scaffolds and a N50 of 42,72 Mb.

### Brain dissection, cell count and PGLS analysis

Fish were anesthetized in a tricaine methanesulfonate (MS-222) solution (500 mg/l) and their body mass and length were measured. After additional intramuscular injection of a ketamine/xylazine mixture (3:1), they were perfused transcardially with wormed phosphate buffered saline containing 0.1% heparin followed by cold 4% paraformaldehyde solution. The dorsal part of the skull was largely removed and the head with exposed brain was fixed in the 4% paraformaldehyde for about 30-60 minutes. Brain was then dissected and weighed using a Mettler Toledo MX5 microbalance (Mettler Toledo, Columbus, Ohio). Brains were postfixed for additional 2-3 days, transferred to an antifreeze solution (30% glycerol, 30% ethylene glycol, 40% phosphate buffer) and kept frozen at 20°C until processing.

Olfactory bulbs were separated from the rest of brain. The two parts were weighed and the total number of their constituent cells was determined following the procedure of isotropic fractionator^45^. Briefly, each brain division was homogenized in a dissociation solution (40 mM sodium citrate solution with 1% Triton X-1000) using glass tissue grinders (0.5 ml or 1 ml, Ningbo Ja-Hely Technology Co., Ltd., China). When turned into an isotropic suspension of free cell nuclei, homogenates were stained with the fluorescent DNA marker 4′,6-Diamidino-2-phenylindole dihydrochloride (DAPI) (Sigma-Aldrich), its volume was determined using an Eppendorf Xplorer 5–1000 μL electronic pipette (Eppendorf, Hamburg, Germany) and kept homogenous by agitation. The total number of nuclei in suspension, and therefore the total number of cells in original tissue, was estimated by determining the number of nuclei in 10 μl samples drawn from the homogenate (Supplementary Table 7). At least six aliquots were sampled and counted using a Neubauer improved counting chamber (BDH, Dagenham, Essex, UK) at the AxioImager.A2 microscope (Carl Zeiss AG, Jena, Germany) equipped with epifluorescence and appropriate filter settings; additional aliquots were assessed when needed to reach the coefficient of variation among counts ≤ 0.1.

To assess the relationship between the number of olfactory genes and the number of olfactory bulb cells, we performed phylogenetic least squares regression using the gls function in the R package nlme^46^ (R package version 3.1-159, https://CRAN.R-project.org/package=nlme). The response variable was the number of cells in the olfactory bulbs and the explanatory variables were the number of cells in the rest of the brain (to control for overall brain size) and the number of OLF genes. All variables were log10-transformed prior to analysis. To visualize the relationship, the residuals from regression of olfactory bulb cells on the rest of brain cells were plotted against the number of OLF genes.

## Acknowledgments

Computational resources were supplied by the project “e-Infrastruktura CZ” (e-INFRA CZ LM2018140) supported by the Ministry of Education, Youth and Sports of the Czech Republic. We also thank the Long Read Team of the DRESDEN Concept Genome Center, part of the MPI-CBG and the technology platform of the CMCB at the TU Dresden. We thank Vojtech Kašpar, Lucie Marhounová and Martin Kocourek for helping with fish sampling. We would like to thank Miriam Martínez de Bustos for drawing most fish silhouettes, and Jan Havlíček, Enrique Navas, Manuel Irimia, Isabel Almudi and Nacho Maeso for reading and commenting the manuscript. This project was supported by the PRIMUS Research Programme (Charles University), the Czech Science Foundation (21-31712S, 20-28135S), and Swiss National Science Foundation (PROMYS – 166550), DB was further funded by Fond Junior (Post-doc) and Mobility (both Charles University). FD research was supported by GAUK project (1108219), SW by DFG (INST 269/768-1) and AN by HFSP LTF LT000400/2019.

## Author contributions

DB and ZM conceived the study. DB performed OLF gene annotation and associated genomic analysis, olfactory organ dissections and RNA extractions, transcriptomic analyses, time-calibrated phylogenetic analysis, and was responsible for the graphical illustrations. FD, YZ, KK and PN carried out brain perfusions and dissections. FD performed cell counting. MS and AN collected and provided the data set on fish nocturnality. KK performed the statistical analyses of ecological traits and PGLS. SW, TB, MP and EM sequenced and assembled the eel catfish genome. DB and ZM wrote the manuscript draft, and all co-authors commented and approved the final version.

## Data and materials availability

Eel catfish genome assembly and associated raw data can be found with the following accession codes: Primary assembly: BioProject accession PRJNA834615, Genome accession JAMASW000000000. Alternate assembly: BioProject accession PRJNA834614, Genome accession JAMASX000000000. Biosample: SAMN28052917. PacBio HiFi data: SRA accession SRR19049190. HiC data: SRA accession SRR19049189. All fastq files of the RNA-seq samples generated for this project are available at SRA within BioProject PRJNA899054.

## Supplementary Figures

**Supplementary Fig. 1:**
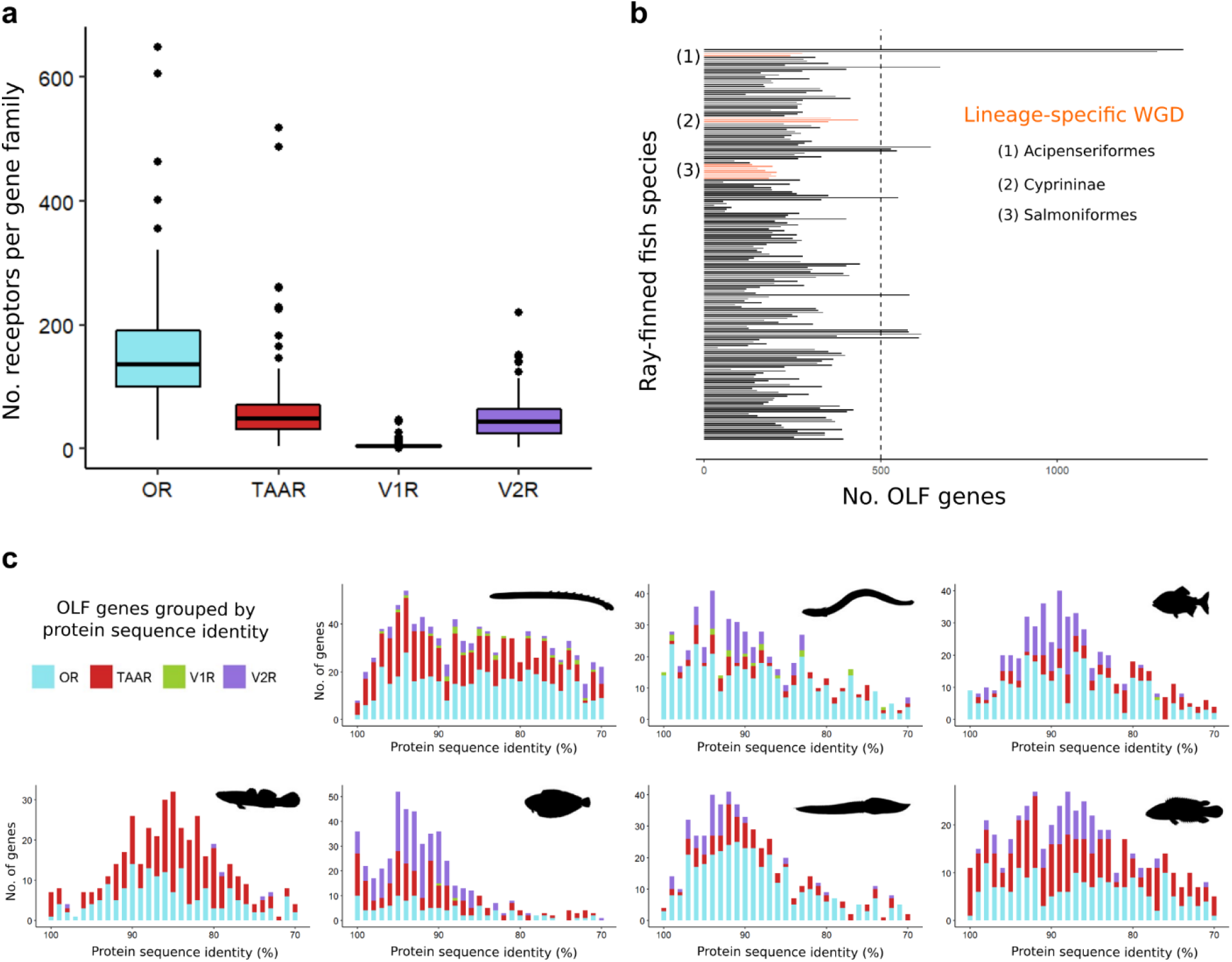
Evolutionary dynamics of OLF gene repertoires. (**a**) Comparison of the distribution in terms of receptor numbers between OLF gene families in our dataset of 201 fish species. Median and average values are as follow, respectively: OR (136,152), TAAR (49,61), V1R (5,6) and V2R (44,47). (**b**) Fish clades affected by lineage-specific whole genome duplications (WGD) other than the teleost-specific 3R are highlighted in orange. None of them belong to the orders containing more than 500 OLF genes. (**c**) Number of receptor genes showing a decreasing percentage of sequence similarity at the protein level relative to other genes for seven species with the highest OLF repertoires in Fig. 1. Only gene groups between 100% and 70% of identity are plotted for visualization purposes. While species have a small relative number of identical receptors (100% identity), differences in the duplication dynamics among families are observed.

**Supplementary Fig. 2:**
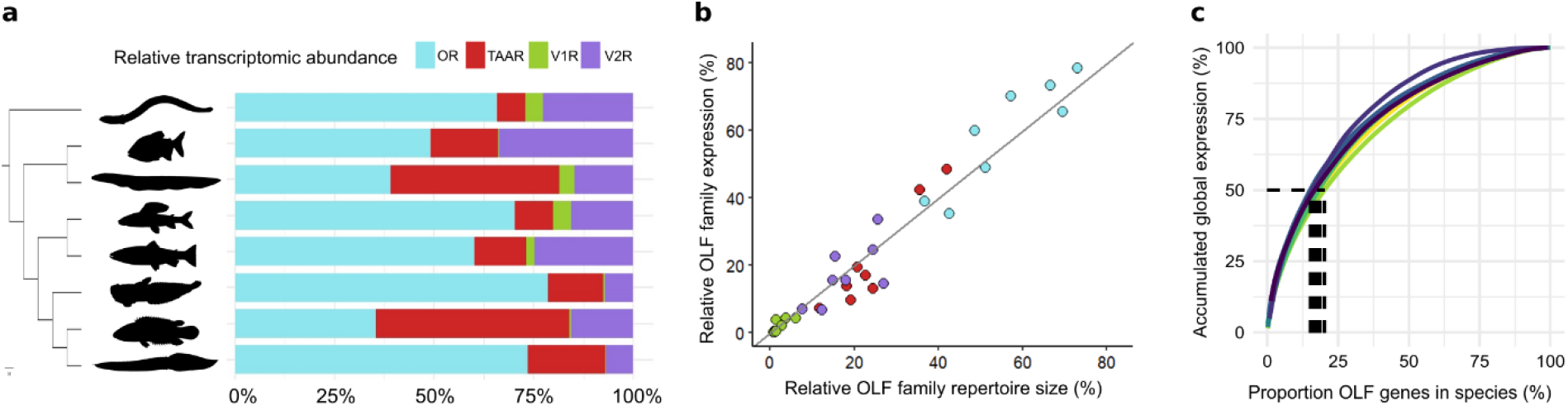
Evolution of OLF genes transcriptomic abundance in eight ray-finned fish species. (**a**) Relative transcriptomic abundance of each OLF family in the olfactory tissue of eight sequenced species. Species in the plot are, from top to bottom: European eel (*Anguilla anguilla*, 669 genes), red-bellied piranha (*Pygocentrus nattereri*, 530 genes), electric eel (*Electrophorus electricus*, 332 genes), grayling (*Thymallus thymallus*, 85 genes), brown trout (*Salmo trutta*, 183 genes), amazon toadfish (*Thalassophryne amazonica*, 141 genes), climbing perch (*Anabas testudineus*, 615 genes) and zig-zag eel (*Mastacembelus armatus*, 608 genes). (**b**) Relation between relative repertoire size and its transcriptomic abundance is shown for each family from the eight studied species. Gray line indicates a hypothetical perfect correspondence. (**c**) Proportion of individual receptors accumulating half of the total OLF gene expression is indicated with dashed lines for each species. Values range from around 15% in the brown trout to approximately 20% in the climbing perch.

**Supplementary Fig. 3:**
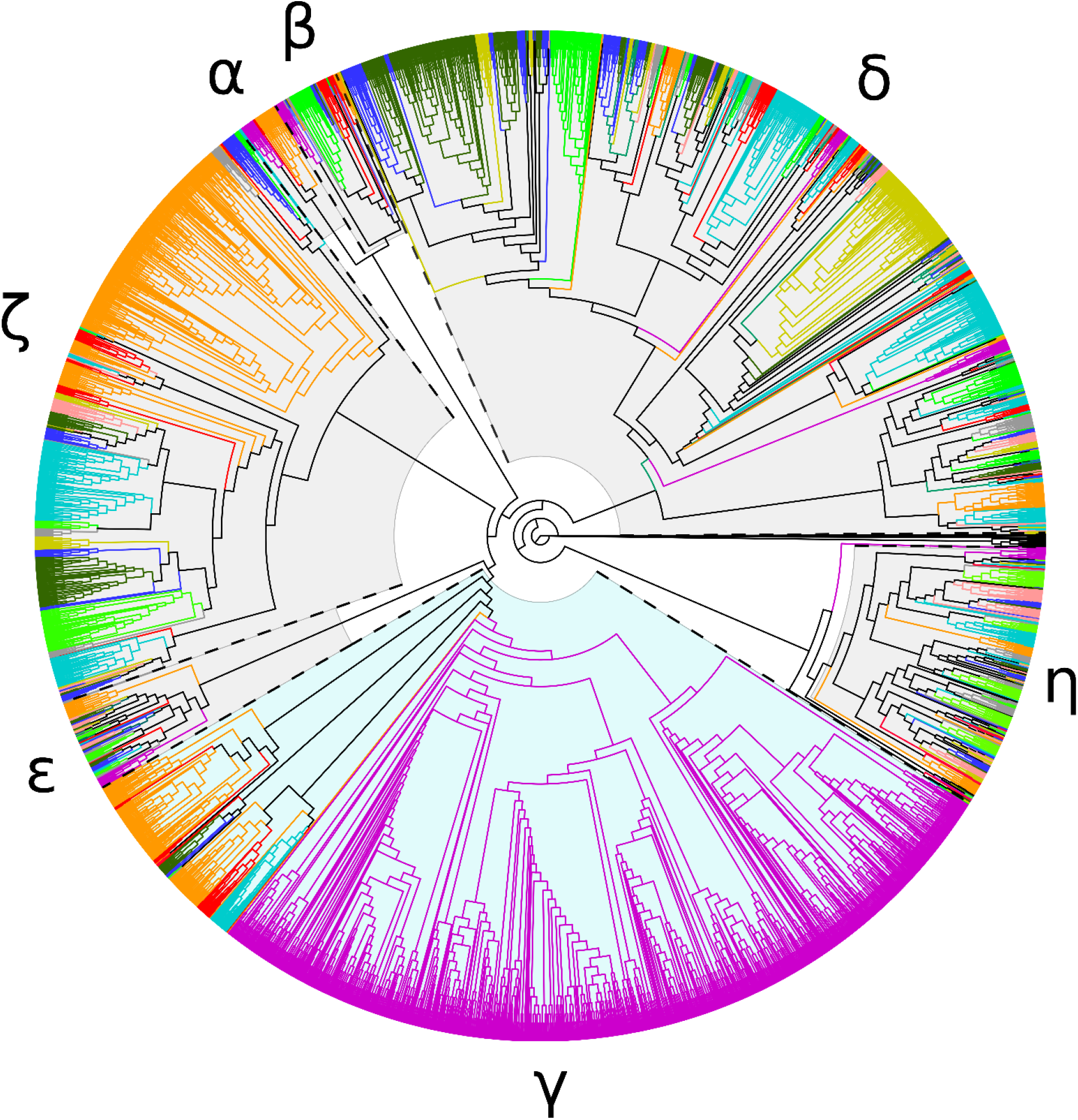
Phylogenetic analysis of OR genes subtypes in ray-finned fishes with large OLF repertoires. Maximum-likelihood phylogenetic tree containing OR protein sequences from those species with more than 500 OLF genes highlighted in Fig. 1, plus spotted gar, zebrafish, and three tetrapods (*Gallus gallus, Anolis carolinensis* and *Xenopus tropicalis*) obtained from public sources. Length of branches is transformed, with tips aligned to help visualization. Distinct branch colors are used for genes from each species or lineage. Color identities are as follow: tetrapod species in light purple, ropefish (*Erpetoichthys calabaricus*) in orange, spotted gar (*Lepisosteus oculatus*) in red, European eel (*Anguilla anguilla*) in aquamarine, tambaqui (*Colossoma macropomum*) in light green, zebrafish (Danio rerio) in grey, four-eyed sleeper (*Bostrychus sinensis*) in dark yellow, Senegalese sole (*Solea senegalensis*) in pink, climbing perch (*Anabas testudineus*) in dark blue and zig-zag eel (*Mastacembelus armatus*) in dark green. OR subtypes are indicated with their corresponding greek letter in the outer part: alpha (α), beta (β), gamma (γ), delta (δ), epsilon(ε), zêta (ζ), êta (η). γ-OR receptors are highlighted in light blue background. Sequences were aligned with MAFFT and IQTREE2 was used for phylogenetic reconstruction.

**Supplementary Fig. 4:**
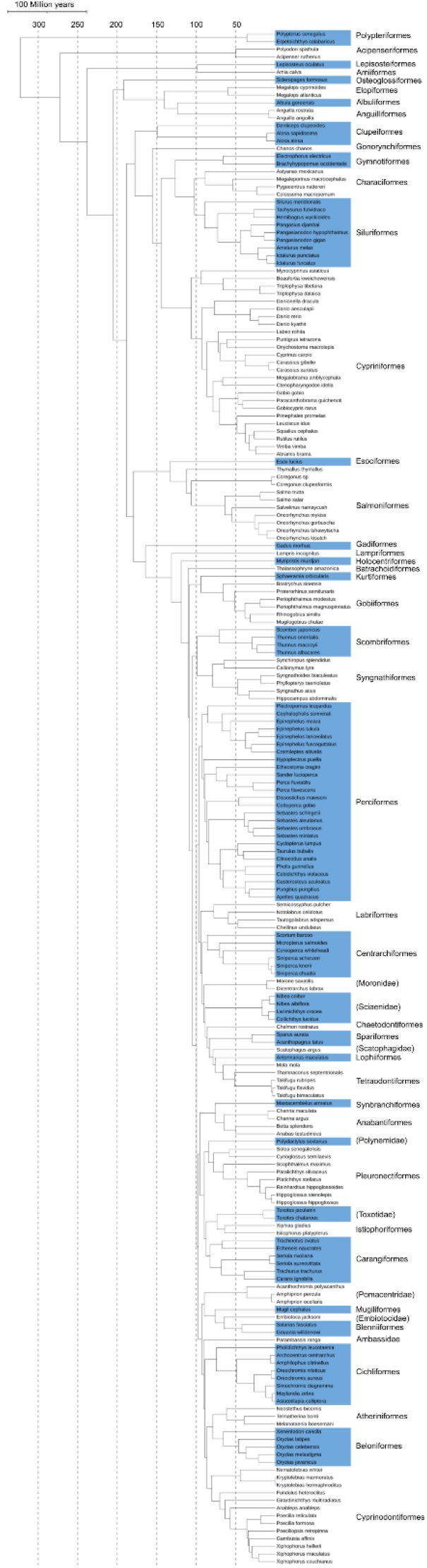
Time-calibrated phylogenetic tree of studied ray-finned fish species. The tree shows phylogenetic position and evolutionary distance in millions of years between 201 species. Blue and white alternated backgrounds are used to separate clades. Taxonomic orders are indicated except for lineages belonging to order-level *incertae sedis*, where taxonomic family is in brackets.

## Supplementary Tables and Data captions

**Supplementary Table 1**: **Number of OLF genes identified in genome assemblies**. Table showing the number of genes detected for each OLF family in a dataset that includes 201 ray-finned fish species. The code name of each assembly and associated values regarding quality measurements are also specified.

**Supplementary Table 2**: **Gene cluster homologies in species with expanded OLF repertoires**. Table reporting homologies between OLF gene clusters for the species shown in Figure 1a and 1b based on syntenic genes. Gene clusters sharing most of the indicated syntenic genes in different species were considered homologous. Homologous origin within species due to duplication or genomic rearrangement are also accounted. Genomic coordinates are indicated for all clusters.

**Supplementary Table 3: Information of transcriptomic samples**. Table listing the RNA-seq data provided and analysed in this study. Information regarding the number of raw, filtered, and mapped reads to OLF genes is also included.

**Supplementary Table 4: Ecological traits in analysed ray-finned fish species**. Table containing discrete classifications of several parameters related to ecological, life-history and behavioural traits for our species dataset. Related factors (same colour) are mutually exclusive, except for salinity in case of species found in different environments.

**Supplementary Table 5: Classification of diurnal and nocturnal activity in fishes**. Table containing diurnal/arrhythmic vs nocturnal activity classification based on the type of evidence. The level of confidence dependent on found references is also indicated for each species.

**Supplementary Table 6: Statistical comparison of the different ecological, life-history and behavioural traits**.

**Supplementary Table 7: Number of cells in the olfactory bulb and brain of twenty-four fish species**. Table showing the estimated number of cells observed in processed brains and olfactory bulbs with the isotropic fractionator. Data for both individual replicates and species is included.

**Supplementary Data 1: Protein sequences and genomic coordinates of OLF genes**. FASTA and General Feature Format (GFF) files containing the protein sequences and genomic coordinates for all identified OLF genes in this study.

**Supplementary Data 2: Alignment of homologous exons for phylogenetic inference**. Multiple alignment in FASTA format containing 640 concatenated homologous exons from the species in our dataset. This alignment was used to build a time-calibrated tree used for phylogenetic corrections in statistical analyses.

## References

1. Poncelet, G. & Shimeld, S. M. The evolutionary origins of the vertebrate olfactory system. Open Biol. 10, 200330 (2022).

2. Niimura, Y. On the Origin and Evolution of Vertebrate Olfactory Receptor Genes: Comparative Genome Analysis Among 23 Chordate Species. Genome Biol. Evol. 1, 34–44 (2009).

3. Silva, L. & Antunes, A. Vomeronasal Receptors in Vertebrates and the Evolution of Pheromone Detection. Annu. Rev. Anim. Biosci. 5, 353–370 (2017).

4. Eyun, S.-I., Moriyama, H., Hoffmann, F. G. & Moriyama, E. N. Molecular Evolution and Functional Divergence of Trace Amine-Associated Receptors. PLoS One 11, e0151023 (2016).

5. Niimura, Y. Olfactory receptor multigene family in vertebrates: from the viewpoint of evolutionary genomics. Curr. Genomics 13, 103–114 (2012).

6. Lin, Q. et al. The seahorse genome and the evolution of its specialized morphology. Nature 540, 395–399 (2016).

7. Policarpo, M. et al. Evolutionary Dynamics of the OR Gene Repertoire in Teleost Fishes: Evidence of an Association with Changes in Olfactory Epithelium Shape. Mol. Biol. Evol. 38, 3742–3753 (2021).

8. Swaney, W. T. & Keverne, E. B. The evolution of pheromonal communication. Behav. Brain Res. (2009). doi:10.1016/j.bbr.2008.09.039

9. Imai, T. & Sakano, H. Odorant receptor gene choice and axonal projection in the mouse olfactory system. Results Probl. Cell Differ. (2009). doi:10.1007/400_2008_3

10. Dang, P., Fisher, S. A., Stefanik, D., Kim, J. & Raper, J. A. Coordination of olfactory receptor choice with guidance receptor expression and function in olfactory sensory neurons. PLoS Genet. (2018). doi:10.1371/journal.pgen.1007164

11. Iwaniuk, A. N. 1.18 - Functional Correlates of Brain and Brain Region Sizes in Nonmammalian Vertebrates. in (ed. Kaas, J. H. B. T.-E. of N. S. (Second E.) 335–348 (Academic Press, 2017). doi:https://doi.org/10.1016/B978-0-12-804042-3.00024-5

12. Zelenitsky, D. K., Therrien, F., Ridgely, R. C., McGee, A. R. & Witmer, L. M. Evolution of olfaction in non-avian theropod dinosaurs and birds. Proc. R. Soc. B Biol. Sci. 278, 3625–3634 (2011).

13. Niimura, Y., Matsui, A. & Touhara, K. Extreme expansion of the olfactory receptor gene repertoire in African elephants and evolutionary dynamics of orthologous gene groups in 13 placental mammals. Genome Res. 24, 1485–1496 (2014).

14. Wright, P. A. & Turko, A. J. Amphibious fishes: evolution and phenotypic plasticity. J. Exp. Biol. 219, 2245–2259 (2016).

15. Ord, T. J. & Cooke, G. M. Repeated evolution of amphibious behavior in fish and its implications for the colonization of novel environments. Evolution (N. Y). 70, 1747– 1759 (2016).

16. Sayer, M. D. J. & Davenport, J. Amphibious fish: why do they leave water? Rev. Fish Biol. Fish. 1, 159–181 (1991).

17. Bressman, N. R., Farina, S. C. & Gibb, A. C. Look before you leap: Visual navigation and terrestrial locomotion of the intertidal killifish Fundulus heteroclitus. J. Exp. Zool. Part A Ecol. Genet. Physiol. 325, 57–64 (2016).

18. Bressman, N. R., Simms, M., Perlman, B. M. & Ashley-Ross, M. A. Where do fish go when stranded on land? Terrestrial orientation of the mangrove rivulus Kryptolebias marmoratus. J. Fish Biol. 95, 335–344 (2019).

19. Bressman, N. R., Hill, J. E. & Ashley-Ross, M. A. Why did the invasive walking catfish cross the road? Terrestrial chemoreception described for the first time in a fish. J. Fish Biol. 97, 895–907 (2020).

20. Van Wassenbergh, S. et al. Evolution: a catfish that can strike its prey on land. Nature 440, 881 (2006).

21. Van Wassenbergh, S. Kinematics of terrestrial capture of prey by the eel-catfish Channallabes apus. Integr. Comp. Biol. 53, 258–268 (2013).

22. Kverková, K. et al. The evolution of brain neuron numbers in amniotes. Proc. Natl. Acad. Sci. 119, e2121624119 (2022).

23. Weiss, L., Manzini, I. & Hassenklöver, T. Olfaction across the water–air interface in anuran amphibians. Cell Tissue Res. 383, 301–325 (2021).

24. Kishida, T. Olfaction of aquatic amniotes. Cell Tissue Res. 383, 353–365 (2021).

25. Mollo, E. et al. Sensing marine biomolecules: smell, taste, and the evolutionary transition from aquatic to terrestrial life. Frontiers in Chemistry 2, (2014).

26. Nevitt, G. A., Dittman, A. H., Quinn, T. P. & Moody, W. J. J. Evidence for a peripheral olfactory memory in imprinted salmon. Proc. Natl. Acad. Sci. U. S. A. 91, 4288–4292 (1994).

27. Meyer, A. et al. Giant lungfish genome elucidates the conquest of land by vertebrates. Nature 590, 284–289 (2021).

28. Chen, S., Zhou, Y., Chen, Y. & Gu, J. fastp: an ultra-fast all-in-one FASTQ preprocessor. Bioinformatics 34, i884–i890 (2018).

29. Patro, R., Duggal, G., Love, M. I., Irizarry, R. A. & Kingsford, C. Salmon provides fast and bias-aware quantification of transcript expression. Nat. Methods 14, 417–419 (2017).

30. Saraiva, L. R. et al. Molecular and neuronal homology between the olfactory systems of zebrafish and mouse. Sci. Rep. 5, 11487 (2015).

31. Ibarra-Soria, X. et al. Variation in olfactory neuron repertoires is genetically controlled and environmentally modulated. Elife 6, e21476 (2017).

32. Zhang, Z., Sakuma, A., Kuraku, S. & Nikaido, M. Remarkable diversity of vomeronasal type 2 receptor (OlfC) genes of basal ray-finned fish and its evolutionary trajectory in jawed vertebrates. Sci. Rep. 12, 6455 (2022).

33. Matschiner, M., Böhne, A., Ronco, F. & Salzburger, W. The genomic timeline of cichlid fish diversification across continents. Nat. Commun. 11, 5895 (2020).

34. Katoh, K. & Standley, D. M. MAFFT multiple sequence alignment software version 7: Improvements in performance and usability. Mol. Biol. Evol. 30, 772–780 (2013).

35. Minh, B. Q. et al. IQ-TREE 2: New Models and Efficient Methods for Phylogenetic Inference in the Genomic Era. Mol. Biol. Evol. 37, 1530–1534 (2020).

36. Betancur-R, R. et al. The tree of life and a new classification of bony fishes. PLoS Curr. tree life (2013).

37. Bürkner, P.-C. brms: An R Package for Bayesian Multilevel Models Using Stan. J. Stat. Softw. 80, 1–28 (2017).

38. Baid, G. et al. DeepConsensus: Gap-Aware Sequence Transformers for Sequence Correction. bioRxiv 2021.08.31.458403 (2021). doi:10.1101/2021.08.31.458403

39. Altschul, S. F., Gish, W., Miller, W., Myers, E. W. & Lipman, D. J. Basic local alignment search tool. J. Mol. Biol. 215, 403–410 (1990).

40. Cheng, H., Concepcion, G. T., Feng, X., Zhang, H. & Li, H. Haplotype-resolved de novo assembly using phased assembly graphs with hifiasm. Nat. Methods 18, 170–175 (2021).

41. Guan, D. et al. Identifying and removing haplotypic duplication in primary genome assemblies. Bioinformatics 36, 2896–2898 (2020).

42. Zhou, C., McCarthy, S. A. & Durbin, R. YaHS: yet another Hi-C scaffolding tool. bioRxiv 2022.06.09.495093 (2022). doi:10.1101/2022.06.09.495093

43. Poplin, R. et al. A universal SNP and small-indel variant caller using deep neural networks. Nat. Biotechnol. 36, 983–987 (2018).

44. Danecek, P. et al. Twelve years of SAMtools and BCFtools. Gigascience 10, (2021).

45. Herculano-Houzel, S. & Lent, R. Isotropic Fractionator: A Simple, Rapid Method for the Quantification of Total Cell and Neuron Numbers in the Brain. J. Neurosci. 25, 2518 LP – 2521 (2005).

46. Pinheiro, J. C., Bates, D. & DebRoy, S. The R Core Team nlme: Linear and Nonlinear Mixed Effects Models. R Packag. nlme version 3, 1–83 (2007).

